# Human basigin (CD147) does not directly interact with SARS-CoV-2 spike glycoprotein

**DOI:** 10.1101/2021.02.22.432402

**Authors:** Robert J. Ragotte, David Pulido, Francesca R. Donnellan, Giacomo Gorini, Hannah Davies, Juliane Brun, Lloyd D. W. King, Katherine Skinner, Simon J. Draper

## Abstract

Basigin, or CD147, has been reported as a co-receptor used by SARS-CoV-2 to invade host cells. Basigin also has a well-established role in *Plasmodium falciparum* malaria infection of human erythrocytes where it is bound by one of the parasite’s invasion ligands, reticulocyte binding protein homolog 5 (RH5). Here, we sought to validate the claim that the receptor binding domain (RBD) of SARS-CoV-2 spike glycoprotein can form a complex with basigin, using RH5-basigin as a positive control. Using recombinantly expressed proteins, size exclusion chromatography and surface plasmon resonance, we show that neither RBD nor full-length spike glycoprotein bind to recombinant human basigin (either expressed in *E. coli* or mammalian cells). Given the immense interest in SARS-CoV-2 therapeutic targets, we would caution the inclusion of basigin in this list on the basis of its reported direct interaction with SARS-CoV-2 spike glycoprotein.

**Importance:** Reducing the mortality and morbidity associated with COVID-19 remains a global health priority. Critical to these efforts is the identification of host factors that are essential to viral entry and replication. Basigin, or CD147, was previously identified as a possible therapeutic target based on the observation that it may act as a co-receptor for SARS-COV-2, binding to the receptor binding domain of the spike protein. Here, we show that there is no direct interaction between the RBD and basigin, casting doubt on its role as a co-receptor and plausibility as a therapeutic target.

## Introduction

Since the emergence of SARS-CoV-2 as the cause of the ongoing COVID-19 pandemic, there has been a rush to identify therapeutic targets that could reduce the immense human and economic toll of COVID-19. Receptors required for viral entry are a natural consideration for druggable targets, as receptor-blockade could both prevent infection, if a drug is delivered prophylactically, or treat infection in a therapeutic setting by stopping the spread of the virus to other tissues and organs. Moreover, there may be existing monoclonal antibodies (mAbs) approved for clinical use that target these receptors.

After the release of the genome sequence of SARS-CoV-2, the primary entry receptor was rapidly identified as angiotensin converting enzyme 2 (ACE2) (1–5). This is the same entry receptor used by some other coronaviruses, most notably SARS-CoV-1, a highly similar coronavirus that emerged in 2002 (1, 6). Since this initial identification of ACE2, there has been significant discussion in the literature, both peer-reviewed and pre-print, about other co-receptors or co-factors required for entry (5, 7–9). Transmembrane protease, serine 2 (TMPRSS2) is one such co-factor that has been identified and subsequently validated by multiple groups, which cleaves the spike protein to facilitate entry (5, 10, 11).

Another co-receptor that gained some attention is CD147, or basigin, which was first described as a spike glycoprotein co-receptor in the pre-print literature in March 2020 and which has since been published in a peer-reviewed journal (9). The authors suggest that spike binding to basigin has important functional implications for viral entry, making basigin-blockade an attractive therapeutic target (9). Since this initial finding, basigin has been included in discussions of SARS-CoV-2 co-receptors (7, 12–20).

Basigin is ubiquitously expressed in human tissues, and forms a complex with monocarboxylate transporters (MCTs), the glucose transporter GLUT1, integrins α_3_β_1_ and α_3_β_1_, among others (21). In the context of infectious disease, basigin has also been well-characterised as an essential receptor for *Plasmodium falciparum* invasion into human erythrocytes, during which it is bound by the malaria parasite’s reticulocyte binding protein homolog 5 (RH5) (22, 23). Based on the initial observation that appeared to show basigin binds to the SARS-CoV-2 spike glycoprotein receptor binding domain (RBD) (9), clinical trials were initiated investigating an anti-basigin mAb as a therapeutic for COVID-19 (24) (ClinicalTrials.gov identifier NCT04275245).

Having worked extensively with basigin in the context of its RH5 interaction, we aimed to validate the finding that basigin directly interacts with the receptor binding domain of the SARS-CoV-2 spike protein. Here, we show that we could not replicate this finding. Although we see clear binding of recombinant SARS-CoV-2 full-length spike trimer (FL-S) and RBD to ACE2 and the anti-RBD mAb CR3022 (25), we do not see any binding to glycosylated or non-glycosylated basigin through size exclusion chromatography (SEC) or surface plasmon resonance (SPR). Meanwhile, recombinant RH5 shows clear binding to both glycosylated and non-glycosylated basigin through the same methods.

## Materials and Methods

### Recombinant protein expression and purification

A construct for soluble trimeric spike (FL-S) glycoprotein of SARS-CoV-2 (NCBI Reference Sequence: YP_009724390.1), encoding residues M1-P1213 with two sets of mutations that stabilise the protein in a pre-fusion conformation (removal of a furin cleavage site and the introduction of two proline residues: K986P, V987P), was expressed as described previously (26). This construct includes the endogenous viral signal peptide at the N-terminus (residues 1-14), while a C-terminal T4-foldon domain is incorporated to promote association of monomers into trimers to reflect the native transmembrane viral protein. The RBD construct utilised the native SARS-CoV-2 spike signal peptide (1–14) fused directly to residues R319-F541 of the spike glycoprotein which encompasses the binding site for the human receptor ACE2 (26). Both constructs include a C-terminal hexa-histidine (His6) tag for nickel-based affinity purification. FL-S and RBD were transiently expressed in Expi293™ cells (Thermo Fisher Scientific) and protein purified from culture supernatants by immobilised metal affinity followed by gel filtration in Tris-buffered saline (TBS) pH 7.4 buffer.

Full-length SARS-CoV-2 Nucleoprotein (FL-NP, NCBI Reference Sequence: YP_009724397.2, residues M1-A419) was transiently expressed in Expi293™ cells (Thermo Fisher Scientific) intracellularly. The FL-NP construct included a C-tag peptide (EPEA) at the C-terminus (27) for affinity chromatography purification using a C-tag affinity resin (Thermo Fisher Scientific), eluting with 20 mM Tris-HCl, 2 M MgCl_2_ pH 7.4. Affinity purification was followed by size exclusion chromatography in TBS pH 7.4 buffer. Human ACE2 ectodomain (NCBI Reference Sequence: NP_001358344.1, residues Q18-S740) was expressed in Expi293™ cells (Thermo Fisher Scientific) with a preceding murine IgG1 signal peptide, monoFc domain and TEV cleavage site at the N-terminus, and a GTGGS flexible linker and C-tag peptide at the C-terminus. ACE2 containing supernatant was purified by C-tag affinity followed by size exclusion chromatography in TBS pH 7.4 buffer. Human IgG1 CR3022 antibody (28) (GenBank: ABA54613.1 and ABA54614.1) was expressed from heavy and light chain AbVec expression vectors in Expi293™ cells (Thermo Fisher Scientific). CR3022 supernatant was purified using HiTrap Protein G HP (Cytiva) followed by size exclusion chromatography in TBS pH 7.4 buffer. Wild type/native human basigin (BSG-2, residues M1-L206) was expressed in Expi293™ cells (Thermo Fisher Scientific) with C-terminal rat CD4 domains 3+4 (CD4d3+4) tag (to aid expression and solubility) followed by a His6 tag for purification, as described previously (23, 29). Non-glycosylated basigin (also BSG-2)was expressed in *Escherichia coli* with an N-terminal His6 tag followed by a TEV cleavage site, as described previously (22). The recombinant PfRH5 sequence is based on the *P. falciparum* 3D7 clone reference sequence and encodes amino acids E26-Q526. The construct includes a C-terminal C-tag and four mutations to delete N-linked glycosylation sequons (T40A, T216A, T286A and T299A). This construct was expressed as a secreted protein by a stable *Drosophila* S2 cell line (30) and affinity purified using C-tag affinity resin (Thermo Fisher Scientific) followed by a size-exclusion chromatography polishing step in 20 mM Tris, 150 mM NaCl, pH 7.4.

### SEC binding assay

100 μg of recombinant receptor (either mammalian-/*E. coli-*expressed basigin, or ACE2) were mixed in a 1:1 molar ratio with RH5, FL-S, or RBD and then incubated for 1 h at room temperature (RT). After incubation, samples were loaded onto a S200 10/300 column via direct injection using an Äkta Pure (GE Healthcare) and run at 0.8 mL/min at RT. Eluted fractions were collected and run on SDS-PAGE under reducing or non-reducing conditions before staining with Coomassie blue.

### Protein Blots

Samples were diluted 1:4 in Laemelli buffer, with or without dithiothreitol (DTT), and then heated at 95 °C for 5 min before loading onto a pre-cast 4-12 % Bis-Tris polyacrylamide gel (Thermo Fisher Scientific). Samples were run at 200 V for 45 min before staining with Coomassie Blue.

1 μg FL-S or RBD were pipetted onto a 0.2 μm nitrocellulose membrane and allowed to air dry. Immunoblotting was performed using the Invitrogen iBind Western System according to manufacturer’s instructions. CR3022 was used as a primary antibody diluted to 2 μg/mL. Alkaline phosphatase conjugated goat anti-human IgG, Fc-specific, (Sigma) diluted to 1:2000 was used for detection with Sigmafast BCIP/NBT alkaline phosphatase substrate at 1 mg/mL (Sigma-Aldrich).

### Surface plasmon resonance

Basigin, either mammalian- or *E. coli*-expressed, was immobilised on a CM5 chip through amine conjugation using NHS/EDC coupling using a Biacore X100 (GE Healthcare). Samples were run at 30 μL/min with an injection time of 60 s and a dissociation of 200 s. Then 5-step two-fold dilution curves of either RH5, RBD or FL-NP were run over starting at 2 μM, with regeneration of the chip via injection of 10 mM glycine pH 2 for 30 s. Between runs, a single injection of RH5 at 2 μM was carried out to confirm there was no loss in binding activity. For CR3022 binding affinity, approximately 400 response units (RU) of antibody were captured on a protein A chip. Steady-state affinity was determined through an 8-step dilution curve beginning at 1 μM, with 10 mM glycine pH 2 used to regenerate the chip between curves. All curves included one duplicate concentration and were evaluated using the Biacore X100 evaluation software.

### PNGase F treatment

PNGase F treatment was conducted as per manufacturer’s protocol (New England Biolabs). Briefly, 10 μL of basigin at 0.5 μg/μL expressed in either *E. coli* or Expi293™ cells underwent denaturation at 95 °C for 10 min in glycoprotein denaturing buffer (New England Biolabs) followed by immediate cooling on ice for 10 s. Then, the denatured protein was mixed with 2 μL of GlycoBuffer 2, 2 μL 10 % NP-40, 6 μL of water and 1 μL of PNGase F. After incubation for 1 h at 37 °C, samples were analysed by non-reducing SDS-PAGE and stained with Coomassie Blue.

## Results

### Spike and RBD bind human ACE2 via SEC

Initially we produced a panel of recombinant protein reagents. Recombinant human ACE2, SARS-CoV-2 full length spike trimer (FL-S), full-length nucleoprotein (FL-NP), Spike RBD and anti-RBD antibody CR3022 were all expressed by transient transfection in mammalian Expi293™ cells (**Fig. 1A,B**). Glycosylated and non-glycosylated basigin were expressed in Expi293™ and *E. coli* respectively and glycosylation states confirmed by PNGaseF digest (**Fig. S1**). Correct folding of RBD and FL-S was confirmed via dot blot using CR3022, a known SARS-CoV-2 RBD and FL-S binding mAb (28), as the primary antibody (**Fig. 1C**). These data showed all proteins expressed as expected and demonstrated high levels of purity. The FL-S and RBD also showed stability upon freeze-thawing, and retained binding of the conformation-sensitive mAb CR3022 after three freeze-thaw cycles (**Fig. 1C**).

**Figure 1.**
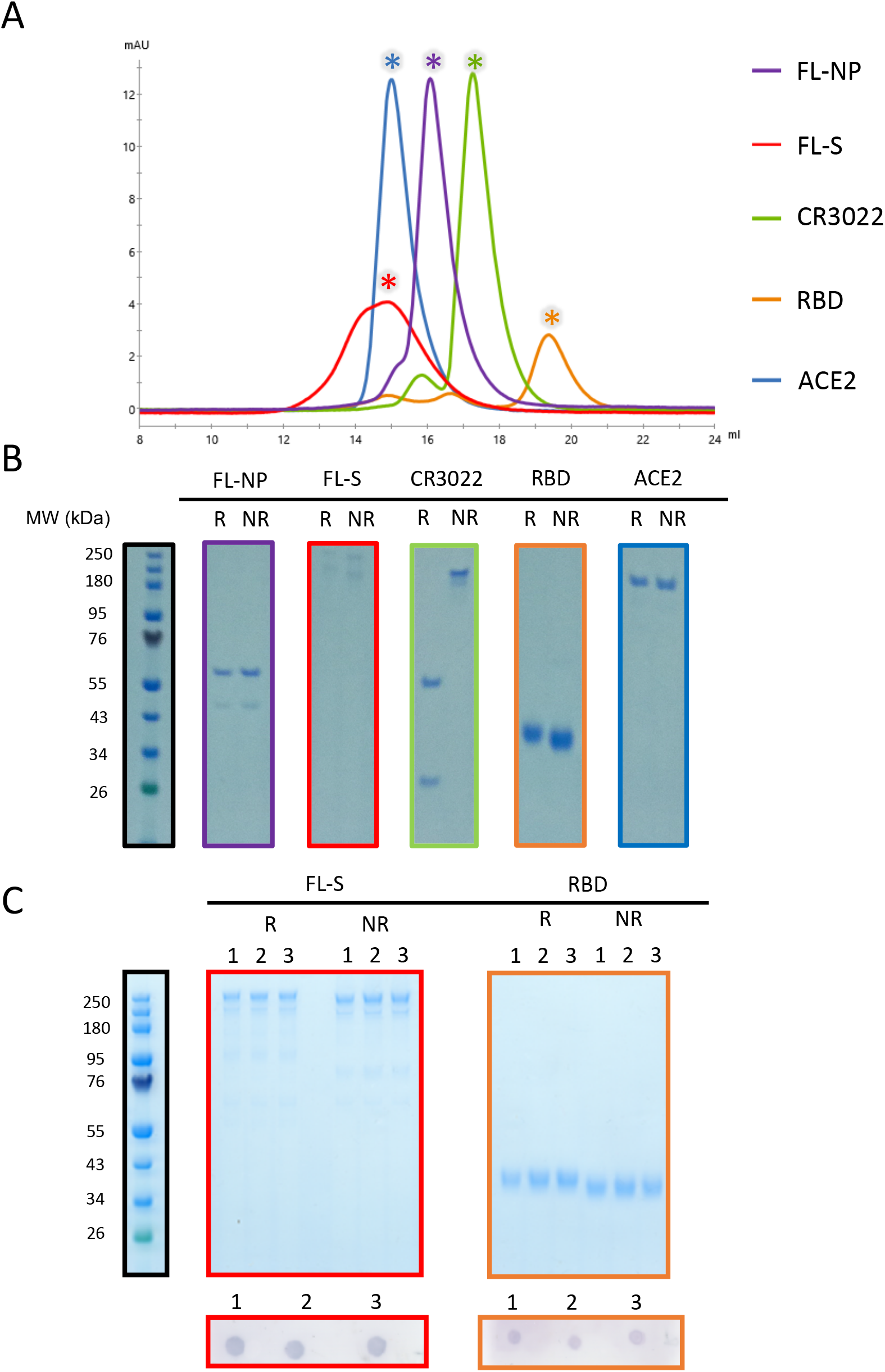
**A**) Size exclusion chromatograms post-purification: 1. FL-NP; 2. FL-S; 3. CR3022; 4. RBD; and 5. ACE2. All proteins were run individually with chromatograms overlaid. Asterisk indicates the fraction run on SDS-PAGE. **B**) Non-reducing (NR) or reducing (R) Coomassie blue-stained SDS-PAGE protein gels of 1 μg of protein from the asterisk indicated fractions. **C**) Freeze-thaw stability of FL-S and RBD. Reducing and non-reducing SDS-PAGE protein gel of 1 μg FL-S (red panel) and RBD (orange panel) after 1, 2 and 3 freeze-thaw cycles. Below each gel a dot-blot is shown, using the CR3022 human mAb on 1 μg FL-S and RBD after 1, 2 and 3 freeze-thaw cycles.

We next confirmed SARS-CoV-2 RBD and FL-S binding to human ACE2 using SEC (**Fig. 2A**). When both RBD and ACE2 were incubated together, the complex eluted at an earlier retention volume as compared to ACE2 alone, indicative of the formation of a higher molecular weight complex. Complex formation was then confirmed using SDS-PAGE whereby both RBD and ACE2 eluted within the same peak at approximately 10 mL, whereas RBD alone normally elutes at approximately 16 mL (**Fig. 2A**).

**Figure 2.**
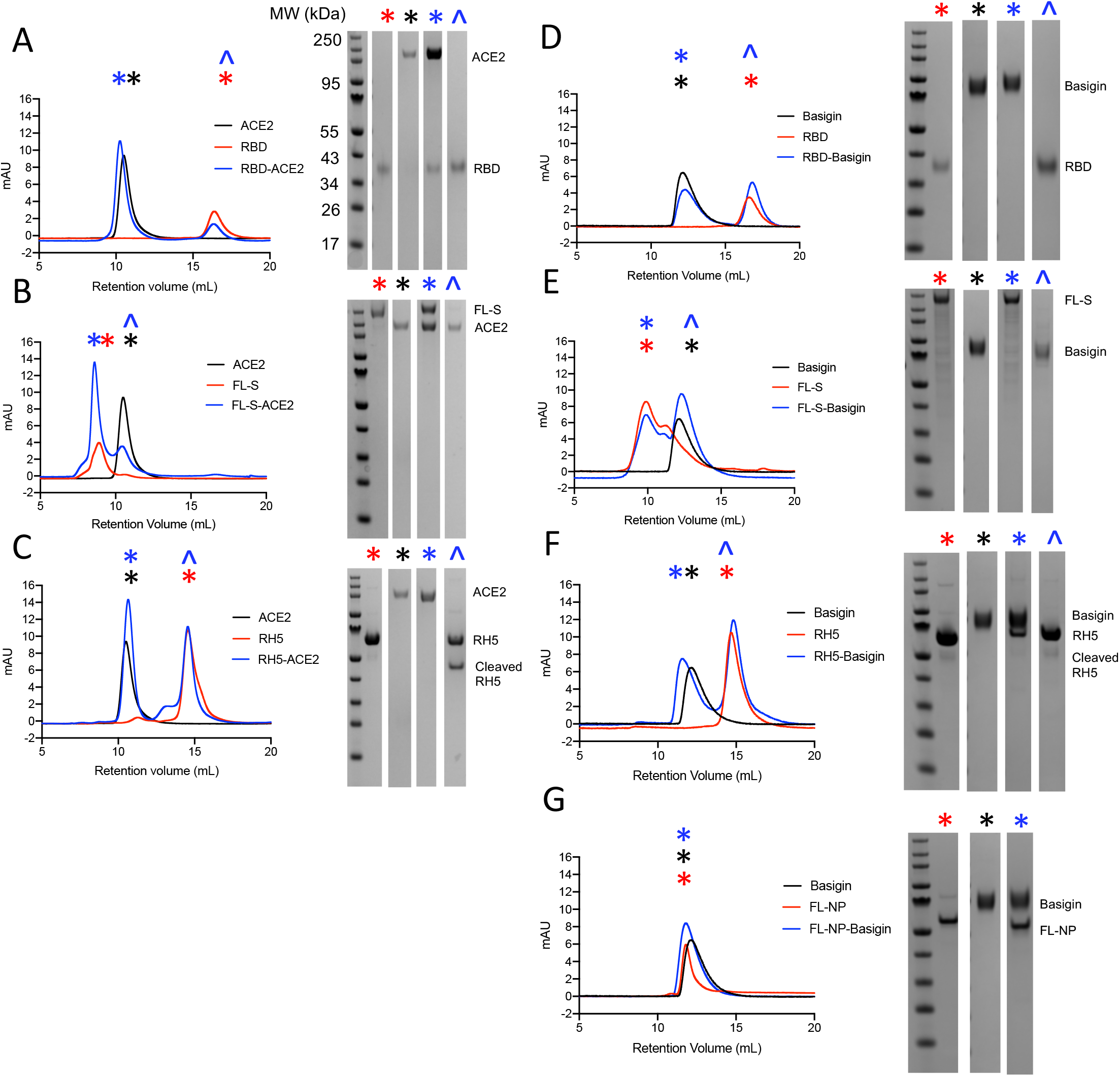
Size exclusion chromatograms (left) and accompanying SDS-PAGE gels (right) assessing complex formation between RH5/RBD/FL-S/RBD and ACE2/Basigin. Symbol on chromatogram indicates which gel corresponds to that peak. Full-length RH5 (~60 kDa) undergoes cleavage at room temperature to yield an ~43 kDa band. **A**) RBD-ACE2; **B**) FL-S-ACE2; **C**) RH5-ACE2; **D**) RBD-Basigin; **E**) FL-S-Basigin; **F**) RH5-Basigin; and **G**) FL-NP-Basigin.

This was next confirmed in the same manner with FL-S trimer, which also eluted as a complex with ACE2 when incubated together, as shown by SDS-PAGE (**Fig. 2B**). Although there is only a small change in retention volume between FL-S alone and the FL-S-ACE2 complex, this can be attributed to the use of an S200 column, whose resolution limits are less than the expected size of the FL-S-ACE2 complex (approximately 680 kDa). Nonetheless, it is clear that the ACE2 eluted with FL-S at approximately 8 mL retention volume, whilst ACE2 alone eluted at approximately 11 mL (**Fig. 2B**). We next demonstrated there was no interaction between the RH5 malaria antigen and ACE2 (as expected), given both proteins eluted at the same retention volume whether alone or mixed together (**Fig. 2C**). These results confirm that our recombinant FL-S, RBD and ACE2 demonstrate the established interactions.

### SARS-CoV-2 spike and RBD do not bind human basigin via SEC

Having confirmed the interaction between FL-S/RBD and ACE2, we proceeded to assess FL-S and RBD binding to glycosylated human basigin using the same methodology. RH5, which acted as the positive control, showed clear binding to basigin, forming a stable complex in solution as confirmed by SEC and SDS-PAGE (**Fig. 2F**). Binding affinity between RH5 and basigin is weaker than the reported values for RBD and basigin (approximately 1μM for RH5 (22, 23) compared to 185 nM for RBD (9)) indicating that this assay should be sufficiently sensitive to detect the RBD-basigin interaction.

SARS-CoV-2 FL-NP was used as a negative control and did not form a complex with basigin. Coincidentally, both FL-NP and basigin elute at the same retention volume, but the absence of any shift to a higher order molecular weight complex when incubated together is consistent with no complex formation (**Fig. 2G**). Next, we observed that there was no detectable binding between either RBD or FL-S and glycosylated basigin, with both RBD and FL-S eluting separately from basigin (**Fig. 2D,E**). Thus, it did not appear that any complex could be formed in solution between these proteins.

Finally, in order to confirm whether glycosylation may affect binding, we performed the experiment again using basigin ectodomain expressed in *E. coli* (**Fig. S1**), as described by Wright *et al.* (22). Again, there was clear binding to RH5, but no discernible binding to either of the FL-S or RBD proteins (**Fig. S2**).

### SARS-CoV-2 spike and RBD do not bind to human basigin via SPR

Although it was clear the reported FL-S/RBD-basigin complex was not stable enough to detect via SEC, we next sought to confirm the previously reported SPR data showing the RBD-basigin interaction (9). To begin, we confirmed that the RBD protein interacted with CR3022 with the expected affinity, via an 8-step dilution curve beginning at 1 μM. The steady state affinity was determined to be 190 nM, consistent with published data on this interaction (25) (**Fig. 3A,B**).

**Figure 3.**
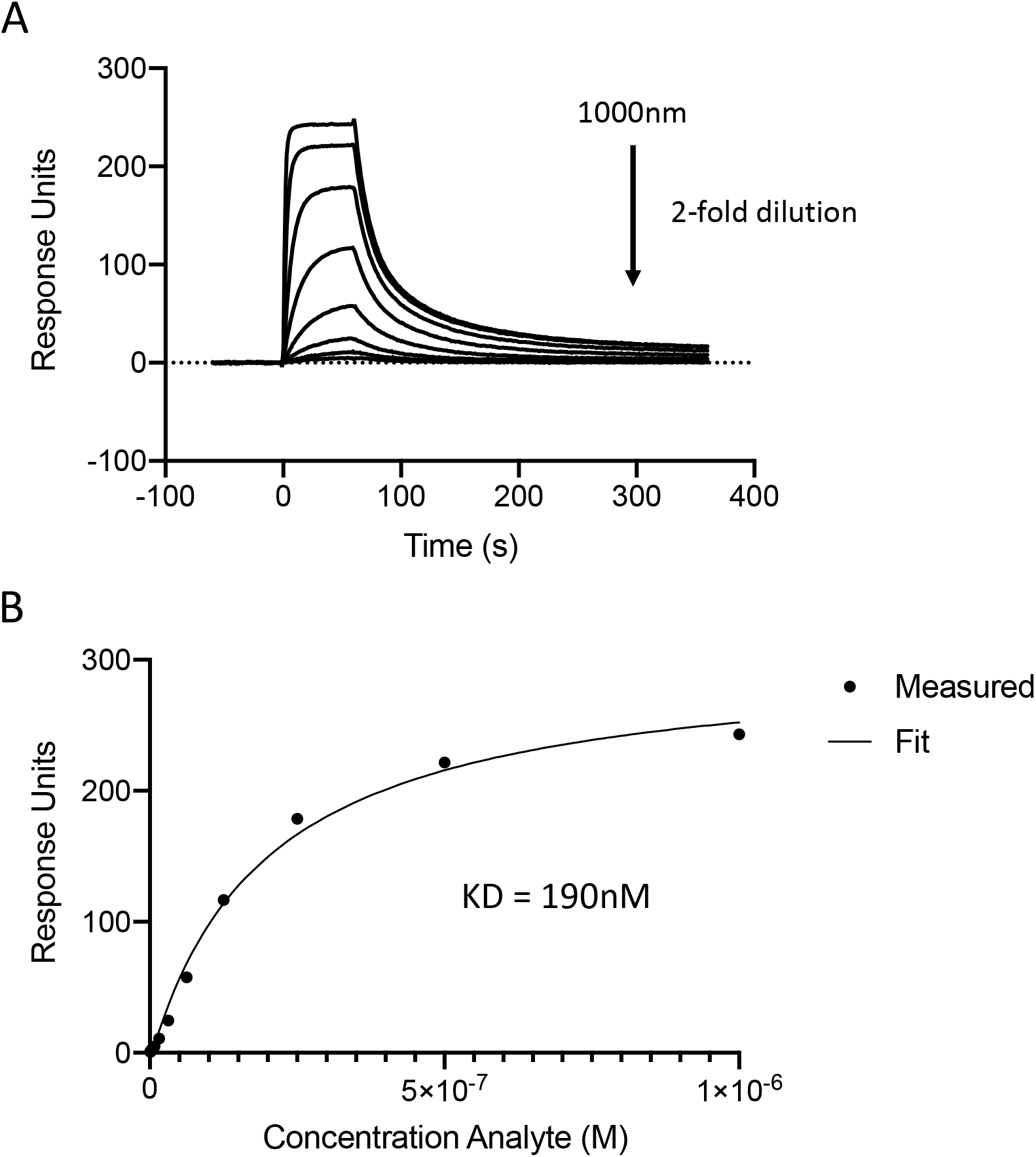
Steady state affinity of CR3022 mAb binding to RBD as assessed using SPR. **A)** Sensorgram of 8-step dilution curve beginning at 1 μM. **B**) Calculation of steady-state affinity.

Next basigin, either glycosylated (**Fig. 4A-C**) or non-glycosylated (**Fig. 4D-F**), was immobilised through amine conjugation on a CM5 chip. RH5, RBD, or FL-NP were then flowed over the chip to determine binding and affinity. RH5 clearly bound to both forms of basigin with a steady-state affinity of approximately 925±16 nM for bacterially-expressed basigin and 665±39 nM for mammalian-expressed basigin, in line with previous reports (22, 23) (**Fig. 4A,D**). However, RBD did not show any discernible binding to either glycosylated (**Fig. 4B**) or non-glycosylated basigin (**Fig. 4E**). FL-NP also did not bind to either form of basigin, as expected (**Fig. 4C,F**).

**Figure 4.**
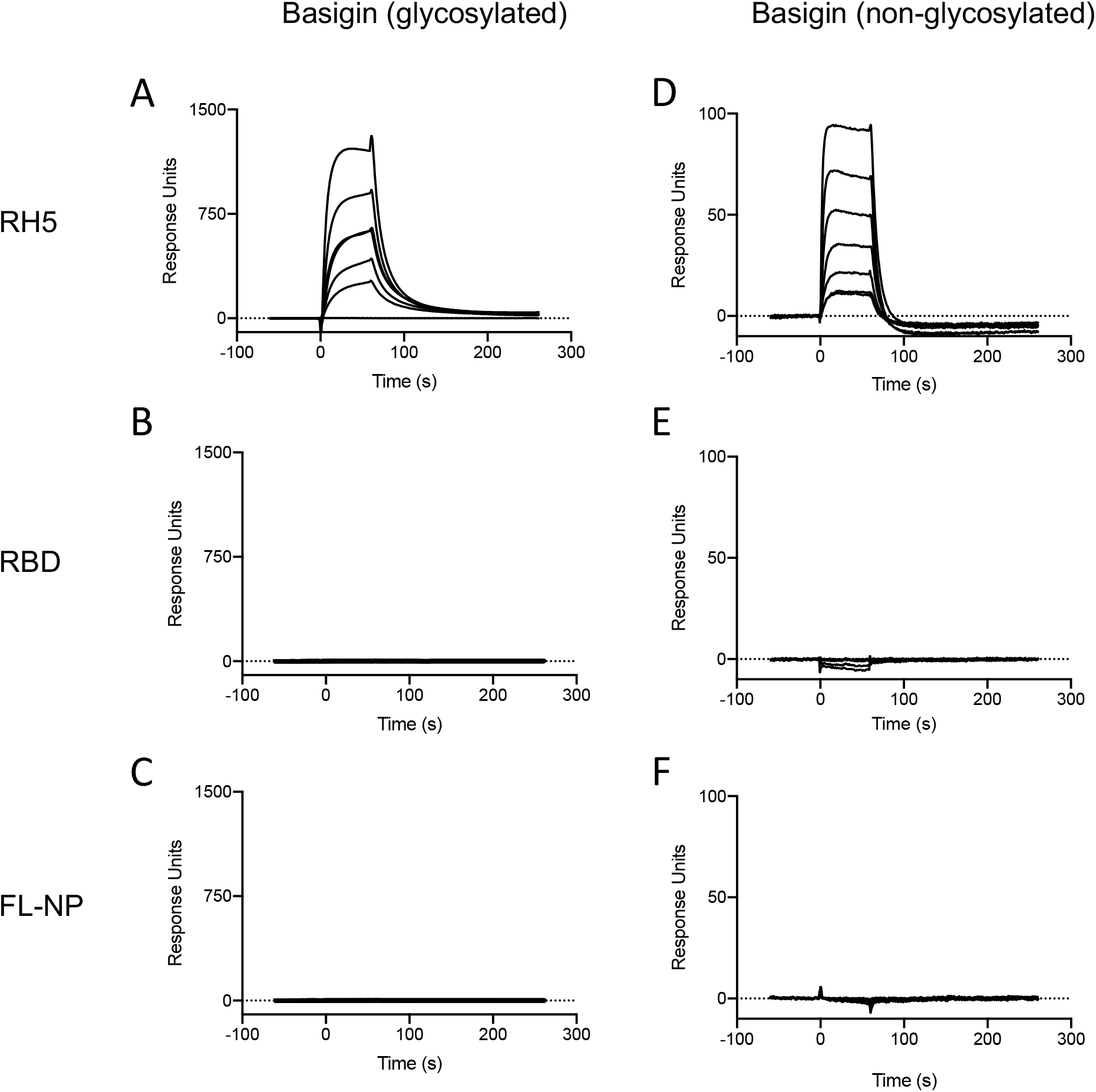
SPR analysis of protein binding interactions. Sensorgrams show binding of each protein to either glycosylated or non-glycosylated basigin (coupled to the chip). Protein binding was assessed along 5-step two-fold dilution curves starting at 2 μM. **A**) Glycosylated basigin binding to RH5; **B**) Glycosylated basigin binding to RBD; **C**) Glycosylated basigin binding to FL-NP; **D**) Non-glycosylated basigin binding to RH5; **E**) Non-glycosylated basigin binding to RBD; **F**) Non-glycosylated basigin binding to FL-NP.

## Discussion

Here, we show that neither SARS-CoV-2 RBD nor full-length spike trimer bind to recombinant human basigin. This is contrary to a previous report in the literature which identified basigin as a co-receptor for SARS-CoV-2 and showed binding of RBD to spike via SPR- and ELISA-based assays (9). The use of anti-basigin mAb in clinical trial has begun on the basis of the original observation that basigin may be required for host cell entry (24). We believe it is necessary to proceed with caution when interpreting the trial data, as further investigation is warranted to determine what role, if any, basigin has in the SARS-CoV-2 invasion process.

Our findings here are also supported by another independent investigation (31). Shilts *et al.* also show evidence that there is no direct interaction between CD147 and full-length spike or its S1 domain using a different set of methods than used here (avidity-based extracellular interaction screening and tetramer-staining of HEK293 cells expressing basigin) (31). Their studies complement the work described here, as they also evaluated this interaction using two different isoforms of basigin and in a cellular invasion assay (31), whereas here we only evaluated the far more abundant basigin-2 isoform (32).

Initial evaluation of meplazumab, an anti-basigin antibody, for treatment of SARS-CoV-2 pneumonia suggested there could be a benefit; however, we interpret these claims with the utmost caution due to the lack of a peer reviewed publication at present and exceedingly small group sizes (24). If indeed these findings hold, it could be due to non-specific anti-inflammatory effect of basigin blockade, as there have been some reports of a pro-inflammatory role of basigin in immune signalling (33–37). Alternatively, other viruses have been reported to utilize CD147 to aid cellular invasion via indirect interactions – including HIV-1 (38) and human cytomegalovirus (HCMV) (39), and these interactions can show cell-type dependence. The *in vitro* effects of meplazumab on SARS-CoV-2 infection and association of this process with endocytosis reported by Wang *et al.* could reflect a similar phenomenon (9); although the basigin knock-down data reported by Shilts *et al.* in the lung epithelial cell line (CaLu-3) would argue against this. Regardless, the benefit of meplazumab is unlikely to be due to the direct inhibition of viral entry, due to the fact that SARS-CoV-2 spike protein does not appear to interact with basigin in our study or that of Shilts *et al* (31)..

Nevertheless, the data surrounding the safety and tolerability of anti-basigin may have implications beyond SARS-CoV-2, as these trials could inform the use of anti-basigin as a malaria prophylactic regardless of its effectiveness in reducing mortality and morbidity due to COVID-19. To date, this has not been pursued in the malaria field beyond *in vitro* assays (40) or humanised mouse models (41), despite demonstration of remarkable potency of anti-basigin mAbs. This is largely due to safety concerns regarding prophylactics that would target a human host protein, as opposed to the parasite, in a vulnerable/infant target population. Should this therapy prove to be safe and well-tolerated, it could be further explored in malaria where basigin has a well-established role in pathogen invasion.

## Conclusion

Recombinant basigin (CD147) does not bind directly to the SARS-CoV-2 RBD. The data presented here do not support the role of basigin as a possible SARS-CoV-2 co-receptor.

## Acknowledgments

The authors are grateful for the assistance of Julie Furze, Amy Boyd, Teresa Lambe (University of Oxford); also to Alain Townsend (University of Oxford) for providing the ACE2 and CR3022 plasmids and Florian Krammer (Icahn School of Medicine at Mount Sinai) for providing the spike and RBD plasmids. The development of SARS-CoV-2 reagents was partially supported by the NIAID Centers of Excellence for Influenza Research and Surveillance (CEIRS) contract HHSN272201400008C.

RJR is funded by the Wellcome Trust Infection, Immunology, and Translational Medicine D. Phil programme, a Rhodes scholarship, and a Canadian Institutes of Health Research Doctoral Foreign Study Award (FRN:157835). FRD holds a UK Medical Research Council (MRC) iCASE PhD studentship (MR/N01796X/1). JB is supported by a Wellcome Trust PhD Programme (203853/Z/16/Z). SJD is a Jenner Investigator, a Lister Institute Research Prize Fellow and a Wellcome Trust Senior Fellow [106917/Z/15/Z]. For the purpose of Open Access, the authors have applied a CC BY public copyright licence to any Author Accepted Manuscript version arising from this submission.

## Author Contributions

RJR, DP, and SJD conceived of the study. RJR, DP and SJD wrote the manuscript. RJR, DP, FRD, GG, HD, JB and LDWK performed experiments. RJR, DP and SJD performed data analysis and interpreted results. KS performed project management. RJR, DP, FRD, GG, HD, JB, LDWK, KS, and SJD reviewed the final manuscript.

## Data and Materials Availability

Requests for materials should be addressed to the corresponding author.

---

**Figure S1.**
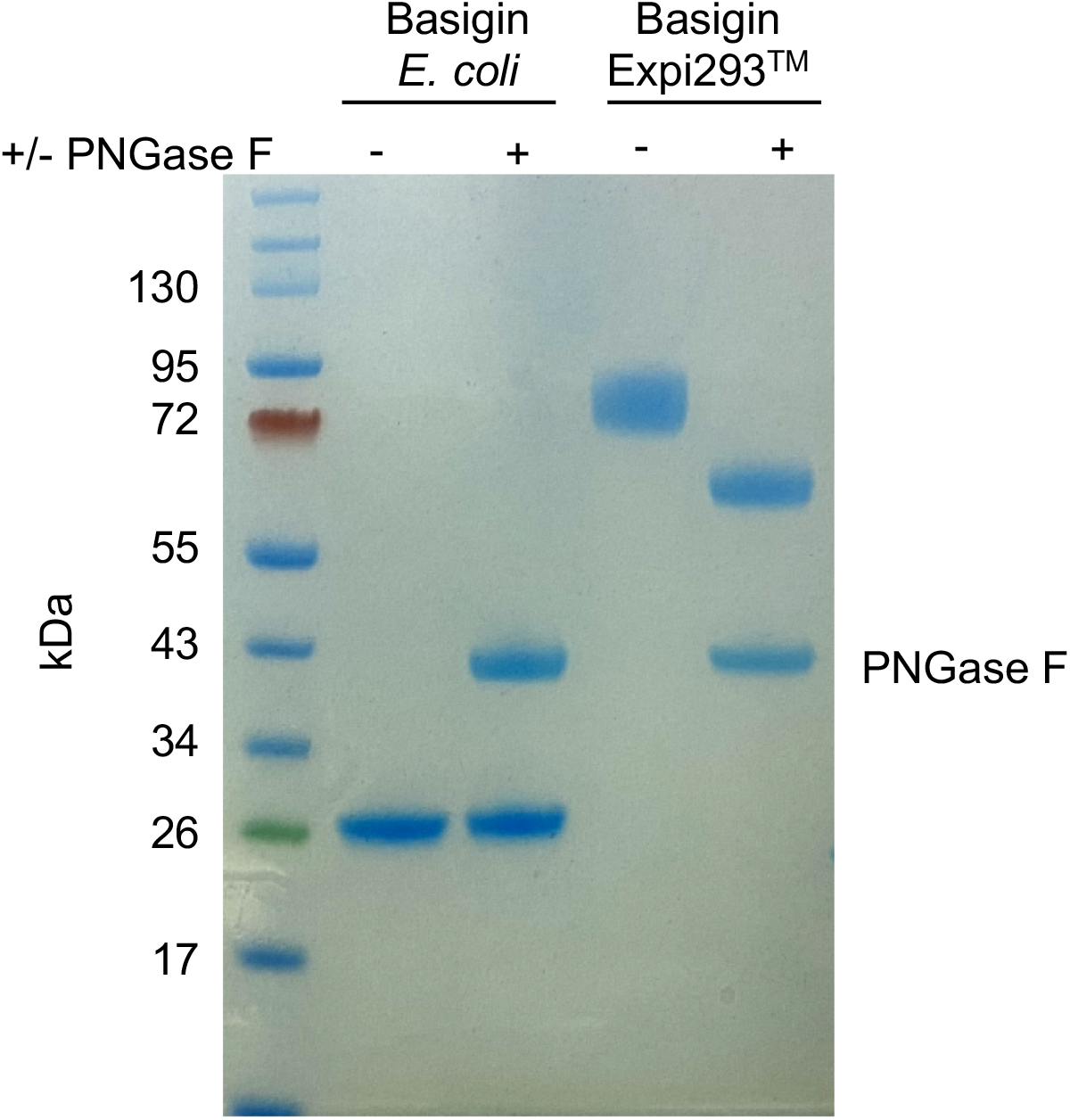
PNGase F digest of *E. coli*-expressed (non-glycosylated) and Expi293™-expressed (glycosylated) basigin. The lower molecular weight of glycosylated basigin after PNGase F treatment is consistent with the loss of glycans. The heavier molecular weight of glycosylated basigin treated with PNGase F compared to non-glycosylated basigin can be attributed to the presence of the rat CD4 domains 3+4 (CD4d3+4) solubility tag (33 kDa).

**Figure S2.**
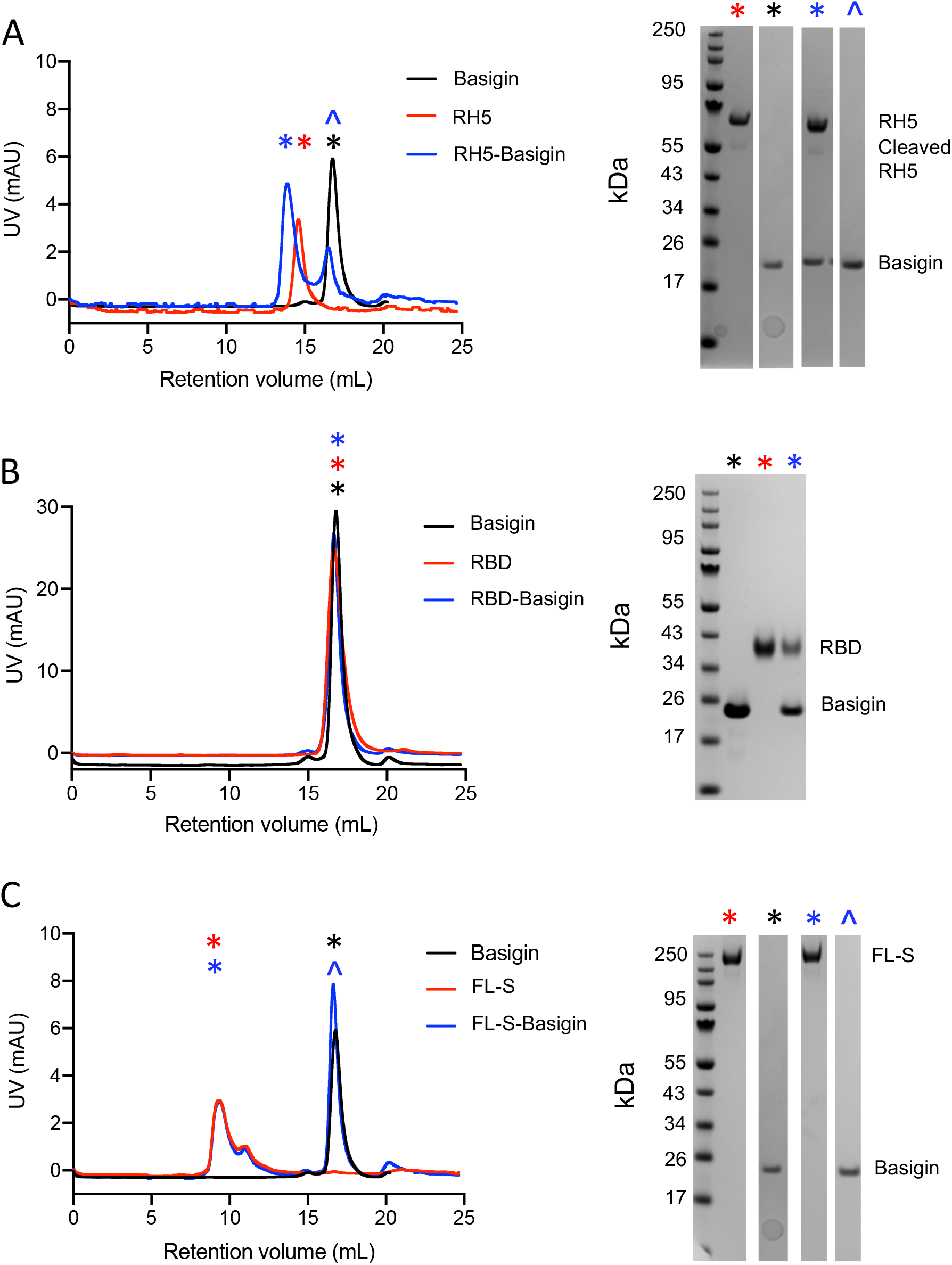
Size exclusion chromatograms (left) and accompanying SDS-PAGE gels (right) of non-glycosylated basigin binding to RH5/RBD/FL-S. **A**) RH5-Basigin; **B**) RBD-Basigin; **C**) FL-S-Basigin.

